# Feature-Based 3D+t Descriptors of Hyperactivated Human Sperm Beat Patterns

**DOI:** 10.1101/2023.04.20.537721

**Authors:** Haydee O. Hernández, Fernando Montoya, Paul Hernández-Herrera, Dan S. Díaz-Guerrero, Jimena Olveres, Alberto Darszon, Boris Escalante-Ramírez, Gabriel Corkidi

## Abstract

The flagellar movement of the mammalian sperm is essential for male fertility as it enables this cell to reach and fertilize an egg. In the female reproductive tract, human spermatozoa undergo a process called capacitation which promotes changes in their motility. Only those spermatozoa that change to hyperactivated (HA) motility are capable of fertilizing the egg; this type of motility is characterized by asymmetric flagellar bends of greater amplitude and lower frequency. Historically, clinical fertilization studies have used two-dimensional analysis to classify sperm motility, although, sperm motility is three-dimensional (3D). Recent studies have described several 3D beating features of sperm flagella, including curvature, torsion, and asymmetries. However, the 3D motility pattern of hyperactivated spermatozoa has not yet been characterized. One of the main difficulties in classifying these patterns in 3D is the lack of a ground-truth reference, as it can be difficult to visually assess differences in flagellar beat patterns. Additionally, only about 10 − 20% of sperm that have been induced to capacitate are truly capacitated (i.e., hyperactivated). In this work, we used an image acquisition system that can acquire, segment, and track sperm flagella in 3D+t. We developed a feature-based vector that describes the spatio-temporal flagellar sperm motility patterns by an envelope of ellipses. Our results demonstrate that the proposed descriptors can effectively be used to distinguish between hyperactivated and non-hyperactivated spermatozoa, providing a tool to characterize the 3D sperm flagellar beat motility patterns without prior training or supervision. We demonstrated the effectiveness of the descriptors by applying them to a dataset of human sperm cells and showing that they can accurately classify the motility patterns of the sperm cells. This work is potentially useful for assessing male fertility or for diagnosing.

## 1 Introduction

Human fertilization requires sperm motility, the ability of the sperm to move and swim through the female reproductive tract to reach the egg. During this journey, the sperm undergoes a process called capacitation, which involves certain biochemical and biophysical changes that enable it to fertilize the egg. Naturally, sperm movement is within a three-dimensional (3D) space in certain regions of the female reproductive tract. However, both research and clinical analysis have been limited to capturing their movement through two-dimensional (2D) images [1–5]. Previous 2D analyses have shown that capacitated sperm exhibit several types of motility [6], including hyperactivated motility, which is characterized by high, asymmetrical flagellar bends, as well as an increase in amplitude and a decrease in the frequency of beating [7, 8]. Only a small percentage (around 10 − 20%) of the millions of ejaculated sperm are truly capacitated showing an hyperactivated motility [9].

The first test used to determine the infertility of a couple is semen analysis, which involves evaluating sperm concentration, motility, morphology, and semen viscosity [10]. In the past, sperm analysis was typically performed visually by a trained technician or scientist using a microscope by tracking the heads of spermatozoa in 2D images, which was often subjective and could produce inconsistent results. To improve accuracy and reduce subjectivity, various classification systems have been developed for this purpose. Computer Assisted Semen Analysis (CASA) has become the gold-standard method for analyzing and classifying sperm motility in 2D, nevertheless, it has been shown that it can produce inconsistent and subjective results as it depends largely on the expertise and training of the user [1, 2]. In recent years, efforts have been made to automate and remove subjectivity from this type of analysis using machine learning and deep learning techniques [6, 11–14]. However, most of these efforts have focused on analyzing sperm head trajectory and CASA parameters extracted from 2D images, leaving the analysis of 3D flagella movement largely unexplored. In the past, imaging the flagella of freely swimming sperm was challenging due to their high beating frequency of up to 25 Hz. Due to the progress of 2D imaging systems, it has been possible to acquire images of the flagellum, for which software systems have been developed to analyze the flagellum in 2D [14], as is the Computer-Assisted Beat-pattern Analysis (CABA) [15]. Even though they are not capable to classify sperm motility even in 2D. The 3D imaging and analysis of sperm flagella kinematics implies several computational challenges, including acquisition, detection, segmentation, tracking, and classification [16], but it provides more information that can help remove subjectivity in the analysis. Despite some recent progress in understanding the 3D flagellar kinematics of single sperm, the classification of 3D flagellar beating patterns in capacitated sperm remains an open research area [14, 16–20].

In this work, we present a new unsupervised classification methodology for 3D flagellar beating patterns. The dataset included both non-capacitated sperm (as a control) and sperm that had been exposed to capacitating conditions, some of which must display hyperactivated motility. There have been different methods developed to classify 3D dynamic patterns [21, 22], but comparing shapes that change over time is a challenging task, especially in the context of motion recognition or video classification [23]. Classifying 3D dynamical patterns is difficult because it requires a deep understanding of the movement and behavior of objects over time and space. There are a variety of techniques that can be used to classify 3D dynamic patterns, including traditional machine learning methods, such as decision trees and support vector machines, as well as more recent techniques based on deep learning, such as convolutional neural net-works (CNNs). These techniques require a large amount of labeled data. Our dataset was unlabeled, because it was not possible to visually identify the different beating patterns by an expert. Additionally, the absence of 3D sperm datasets restricts the capacity to characterize sperm behavior or label 3D sperm data. Hence, we used a different approach for 3D dynamic pattern classification, namely feature extraction, from the 3D+t data followed by an unsupervised classification technique, called hierarchical clustering [24]. We introduce novel descriptors for dynamic motility that are based on a set of ellipses properties enveloping the flagella, which allows to compactly describe the beating information of the flagella from 3D+t. We tested our descriptors using a dataset of 147 free-swimming sperm, experimentally acquired in 3D, that were recorded in a time interval of 1-3 seconds. The dataset was obtained by our group, as described in [16, 18, 25]. We successfully distinguished between non-capacitated and hyperactivated sperm using our proposed dynamic flagella descriptors, which characterize the 3D spatio-temporal flagellar sperm movement patterns by an envelope of ellipses. The shape and similarities between the samples were grouped using hierarchical clustering, resulting in distinct clusters of beating patterns. This is the first time that hyperactivation has been described in 3D, and our study provides the first precedent for future 3D studies.

## 2 Materials and methods

### 2.1 Biological preparations and ethical approval

Under informed written consent and the supervision of the Bioethics committee of the Instituto de Biotecnología, UNAM, young healthy donors supplied human spermatozoa samples by masturbation, after at least 48 hours of sexual abstinence. The semen samples fulfilled the World Health Organization (WHO) requirements [26]. Through a swim up separation, highly motile cells were recovered from the sample that was incubated for 1 hour in Ham’s F-10 medium at 37^°^*C* in a humidified chamber with a 5% CO_2_ concentration. After recovery, cells were centrifuged 5 min at 3000 rpm and half of them were resuspended in a non-capacitating media and the other half in capacitating media. The non-capacitating physiological media consisted of 94 mM NaCl, 4 mM KCl, 2 mM CaCl_2_, 1 mM MgCl_2_, 1 mM Na pyruvate, 25 mM NaHCO_3_, 5 mM glucose, 30 mM Hepes, and 10 mM lactate at pH 7.4. Five *mg/ml* of Bovine Serum Albumin (BSA) and 2 *mg/ml* NaHCO_3_ was added to obtain the capacitating media.

### 2.2 Experimental set-up

The 3*D* + *t* acquisition system consisted of an inverted Olympus IX71 microscope mounted on an optical table [TMC (GMP SA, Switzerland)]. A 60X water immersion objective with *N*.*A*. = 1.00 (Olympus UIS2 LUMPLFLN 60X W) attached to a piezoelectric device P-725 (Physik Instruments, MA, USA) was mounted on the microscope. The piezoelectric along with the objective oscillated with a frequency of 90 Hz and an amplitude of 20 *μm*. A servo-controller E501 via a high current amplifier E-55 (Physik Instruments, MA, USA) was used to fine tune the piezoelectric oscillations. A high speed camera (NAC Q1v) with 8 GB of internal memory was set to record at 8000 fps with a resolution of 640 × 480 pixels. At this speed and resolution, the camera was able to acquire 3.5 seconds. The acquisition system was synchronized via an E-506 function generator (NI USB6211, National Instruments, USA). Temperature of the samples was maintained constant at 37^°^*C* with a thermal controller (Warner Instruments, TCM/CL100).

### 2.3 Dataset

The dataset consisted of 147 human spermatozoa, 60 were exposed to a non-capacitating media and 87 to capacitating media. The data were collected using the system described by Corkidi et al. [16] and the flagellum centerline was reconstructed using the segmentation process described by Hernández-Herrera et al. [18]. From the dataset, it is expected that 10 − 20% of sperm induced to capacitate will display hyperactivated motility [9]. Let 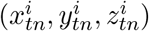 be *n* = 1, 2, …, *N* a set of points of the flagellum’s centerline in a 3D coordinate system at time *t* = 1, 2, …, *T* for each spermatozoa, *i*, in the dataset.

The number of detected flagella and points per single beat may vary over time due to the segmentation process. A complete set of these points for an experiment *i* are shown in Figure 1, each line corresponds to a flagellum reconstruction at a specific time, and the black dots represent the first point in the middle head at each time. The original data was in the laboratory frame of reference. To place all sperm trajectories within the same frame of reference, the flagella were then rotated and translated to align with the *x*−axis starting from the origin, (see Figure 1).

**Figure 1:**
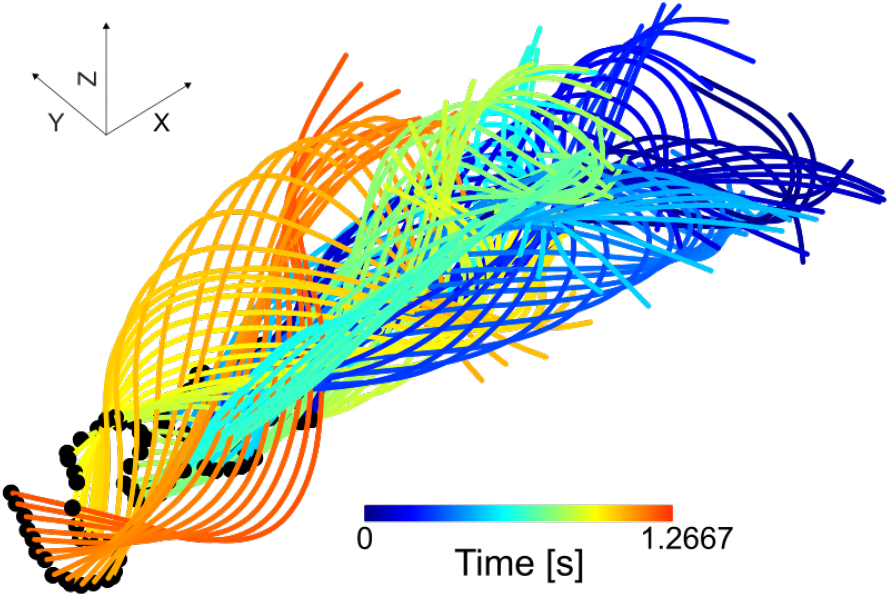
Representation of a tracked and segmented spermatozoon, with the laboratory frame of reference data aligned along the *x*−axis. Each line corresponds to a flagellum reconstruction at a specific time, and the black dots represent the first point of the flagella at each time. The color scale indicates the progression of the beat, with blue representing the initial time advancing towards to red.

### 2.4 Ellipse fitting

We define a flagelloid 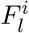 [19] for sperm *i* as the set of points orthogonally projected onto the *Y Z* plane whose *x*-coordinate lie in the interval *I*_*l*_ = [(*l* − 1)· *δ, l* · *δ*) where 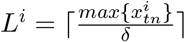 (the symbols ⌈·⌉ denote the ceiling function, e.g., ⌈*π*⌉ = 4) and *δ* is the interval size (as shown in Figure 2)

**Figure 2:**
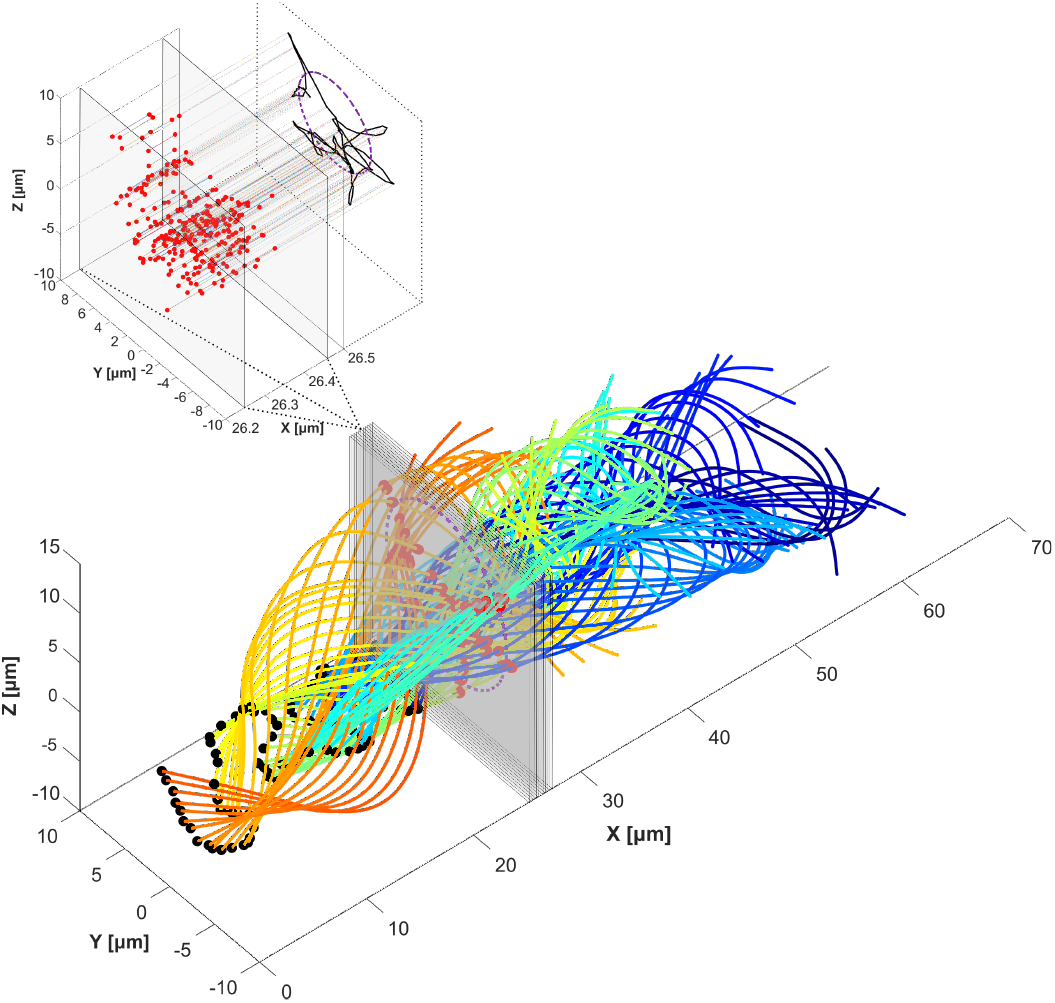
Illustration of the process of fitting an ellipse to a flagelloid. The lower subplot shows the segmented and tracked spermatozoon from Figure 1, with 11 out of 292 gray planes (for visualization purposes only) corresponding to 10 intervals of *δ* each. The upper subplot shows a zoom of a single *δ*− length interval, the red dots represent the flagellum points within that interval. The right upper side displays the orthogonal projection of the red dots, forming a flagelloid (black line) and its fitted ellipse (purple dotted line).

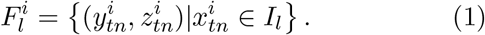

Figure 2 illustrates the fitting of an ellipse (purple) to the points belonging to the flagelloid (Equation 1) using the “Direct fit of least squares of ellipses” method presented in [27]. By performing this process for each flagelloid, a set of transverse ellipses to the *x*−axis are obtained, which describe the motility shape of the sperm (as shown in Figure 3). The interval size, *δ*, is set to ∼ 0.21 *μm*; this value was imposed to ensure that each interval contains at least three points. Ellipses are fitted to the flagelloid only when it has at least this number of points and they are not collinear, to avoid fitting an hyperbola or a parabola (eccentricity, *ε* ≥ 1). The number of ellipses, *E*^*i*^|*E*^*i*^ ≤ *L*^*i*^ are the ellipses that were fitted correctly for spermatozoon *i*.

**Figure 3:**
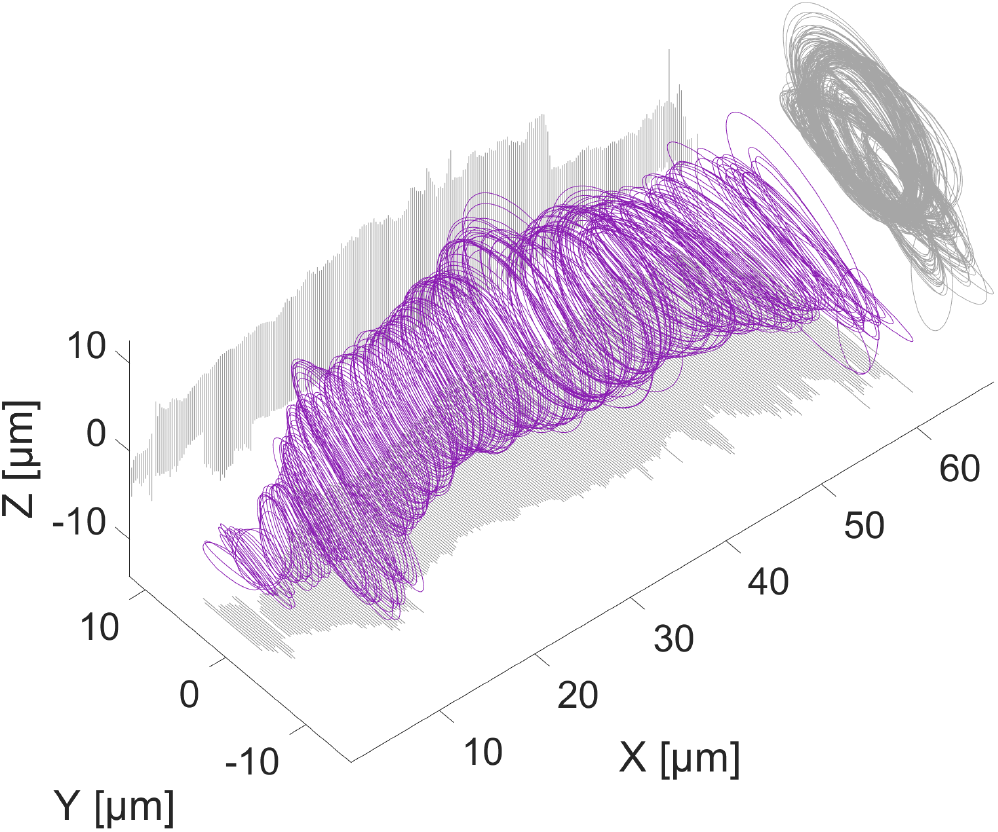
Envelope of ellipses (purple lines) and their XY, XZ and YZ projections (light gray lines) for a spermatozoon. Each ellipse was fitted to an interval, the complete envelope is composed with all the ellipses fitted to the sperm’s beating.

From each ellipse 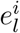, we obtain four parameters:

- the semi-major axis 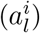
- the semi-minor axis 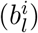
- the angle of the ellipse 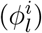 with respect to the *y*−axis
- the eccentricity 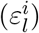.

To describe the variations in the envelope of ellipses, we propose a feature-based vector with the following four components based on an analysis of the obtained data:

- Feature 1 (focus average): 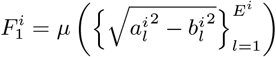
- Feature 2 (angular average): 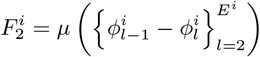
- Feature 3 (eccentricity average): 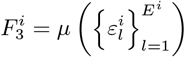
- Feature 4 (semi-major axis deviation): 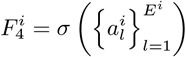

where *μ* is the mean and *σ* is the standard deviation. Feature 1 is the mean distance from the center of each ellipse to its foci, Feature 2 is the mean of the difference between angles, which represents the mean change of angles of the fitted ellipses, Feature 3 is the mean of the semi-major axis and Feature 4 is the standard deviation of the semi-major axis. Then we reduce the dimensionality of the vector using Principal Component Analysis (PCA), resulting in three principal components that account for more than 90% of the variance in the dataset (see Table 1).

**Table 1:** Details of the principal component analysis obtained from the feature-based vector of the envelope of ellipses. The element [*j, k*] is the contribution of the Feature *j* to the *k*-th PC. The dominant features for each principal component are shown in bold.

|  | PC 1 | PC 2 | PC 3 |
| --- | --- | --- | --- |
| Eigenvalue | 2.3862 | 1.0004 | 0.4616 |
| Feature 1 | **0.6142** | 0.0392 | -0.1122 |
| Feature 2 | -0.0242 | **0.9990** | -0.0191 |
| Feature 3 | 0.5712 | -0.0163 | -0.6197 |
| Feature 4 | 0.5440 | 0.0172 | **0.7765** |

### 2.5 Clustering

Our dataset consists of non-capacitated sperm (control) and sperm that have undergone an in-vitro capacitation process, we expect that around 10 − 20% of the capacitated sperm, will be hyperactivated. Since our dataset cannot be labeled, we did not have prior knowledge of the number of clusters in the data, and the ac-tual number of hyperactivated sperm cells is unknown. We choose to apply agglomerative hierarchical clustering, since it does not require knowing a priori how many clusters there are, does not need input parameters, and is less sensitive to outliers compared to other methods. Agglomerative hierarchical clustering [24] is a technique that clusters each of the objects in the dataset based on the dissimilarity metric between them. The clusters are created between pairs of greater similarity and then, they are successively connected with the other pairs of less similarity. This process is represented through a dendrogram that shows the hierarchical relationship between group-pairs in the dataset [24]. We used the Euclidean distance as the dissimilarity metric between the descriptors of each sperm cell with all the others to determine the dissimilarity between each pair of spermatozoa. We used average linkage and “distance” as the criterion to define the clusters based on proximity between the objects. The dendrogram enabled us to visually identify the number of main groups in the dataset, and we identified two main clusters.

## 3 Results

The dissimilarity matrix of the Euclidean distance between each pair of spermatozoa is depicted in Figure 4, illustrating how similar they are based on their feature-based descriptors. The diagonal of the matrix is black since a feature-based descriptor’s distance from itself is 0 (high similarity), while pairs of sperm with higher distances are represented with white (low similarity). The non-capacitated cells and a fraction of the induced to capacitate sperm are found in Cluster 2 (green square and lines in Figure 4). Spermatozoa in Cluster 1 (red squares and lines in Figure 4) have fitted ellipses with semi-major axes that are larger (*max*(*a*) *>* 6.5*μm*) than those in cluster 2 in addition to having *ε* → 1. The ellipse’s eccentricity (*ε*) measures how much it deviates from a circle. This is related to the tendency of the flagellum’s beating pattern, with a more symmetric pattern corresponding to a more circular ellipse (*ε* → 0) and an asymmetric pattern corresponding to a more elongated ellipse (*ε* → 1). Given the previous analysis and considerations, we associate Cluster 2 to the motility pattern of non-capacitated sperm, which includes the control group and a subset of spermatozoa induced to capacitation (those induced that failed-to-capacitate). Cluster 1 is then associated to the hyperactivated beating pattern group, which only includes sperm cells that were induced to capacitate.

**Figure 4:**
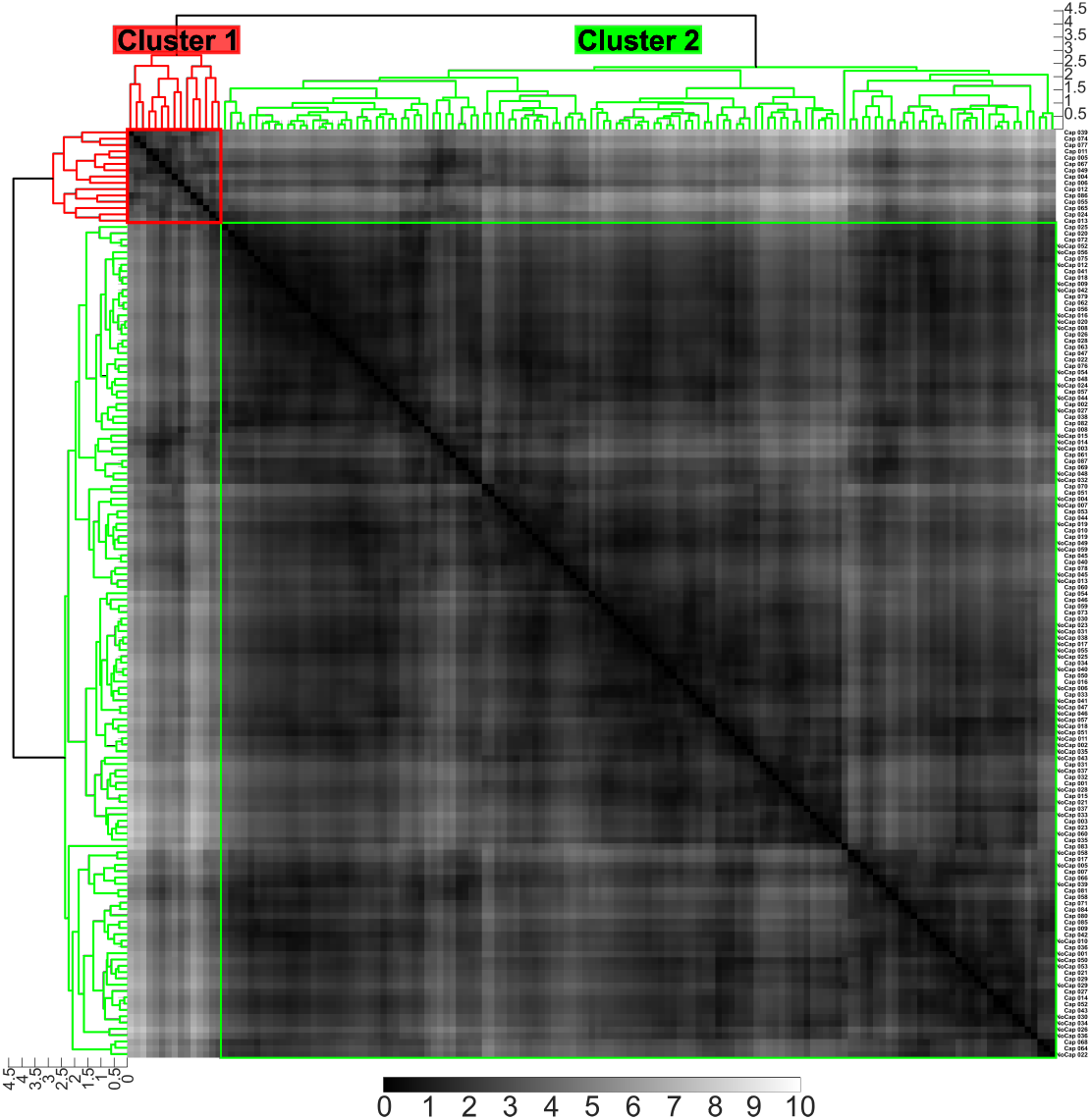
The dissimilarity matrix shows the Euclidean distance between each pair of sperm cells based on their four principal components. The pixel intensity in the matrix corresponds to the distance value. On the right the IDs of each experiment are displayed, where “Cap” refers to the spermatozoa induced to capacitate and “No Cap” belongs to the control group. The dendrogram shows the hierarchical clustering with average linkage; red lines correspond to cluster 1 and green lines to cluster 2.

### 3.1 Validating the classification of the 3D sperm beating patterns

In the literature, it has been described that hyperactivated motility in human spermatozoa is characterized by asymmetric flagellar beating, an increase in amplitude, and a decrease in the frequency of beating [7, 9, 28–32]. To validate our unsupervised classification of motility patterns, we have measured these features (amplitude, frequency, and asymmetry) in 3D+t from our database and proposed some 3D generalizations of the 2D wellknown measures. For this, the first ∼ 2.5 *μm* of arclength of each flagellum was aligned as a head-fixed frame of reference (inner plot from the Figure 5).

**Figure 5:**
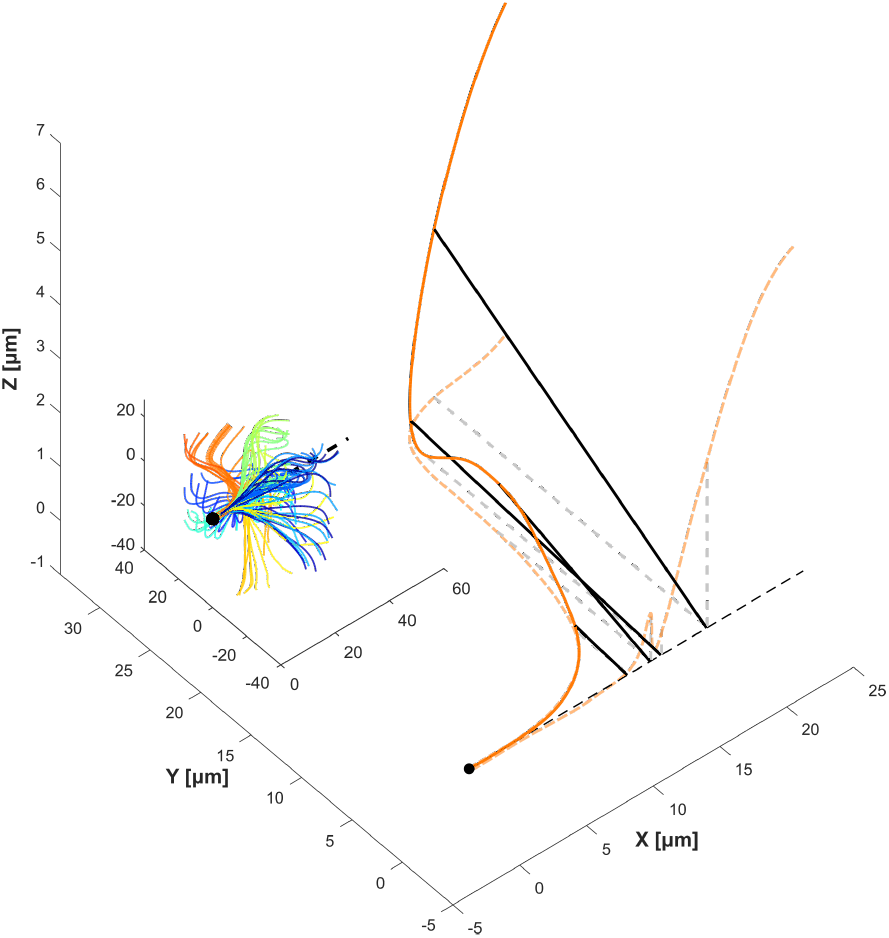
The inside subplot shows the flagellar beating of the spermatozoon aligned and rotated with respect to the first ∼ 2.5 *μm* of arclength, which corresponds to the mid-head of the spermatozoon. The bold orange flagellum is shown in the bottom plot, and the black lines are examples of the amplitude between the points (140, 240, 340 and 390) on the flagellum and the *x*−axis (dotted black line). The gray and orange dotted lines show the projection on the *XY* and *XZ* planes of the flagella, and the distances lines respectively.

We defined the amplitude as the mean distance, denoted by, 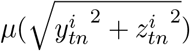 of each point of the flagellum 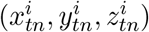 to the *x*−axis for each sperm *i* (Figure 5). The beat frequency is obtained through the analysis of the Fourier transform on each point of the flagellum over the time 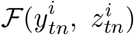. The power spectrum amplitude of each point of the sum of the two components are summed and the frequency with peak amplitude is taken as the maximum frequency (Figure 6):

**Figure 6:**
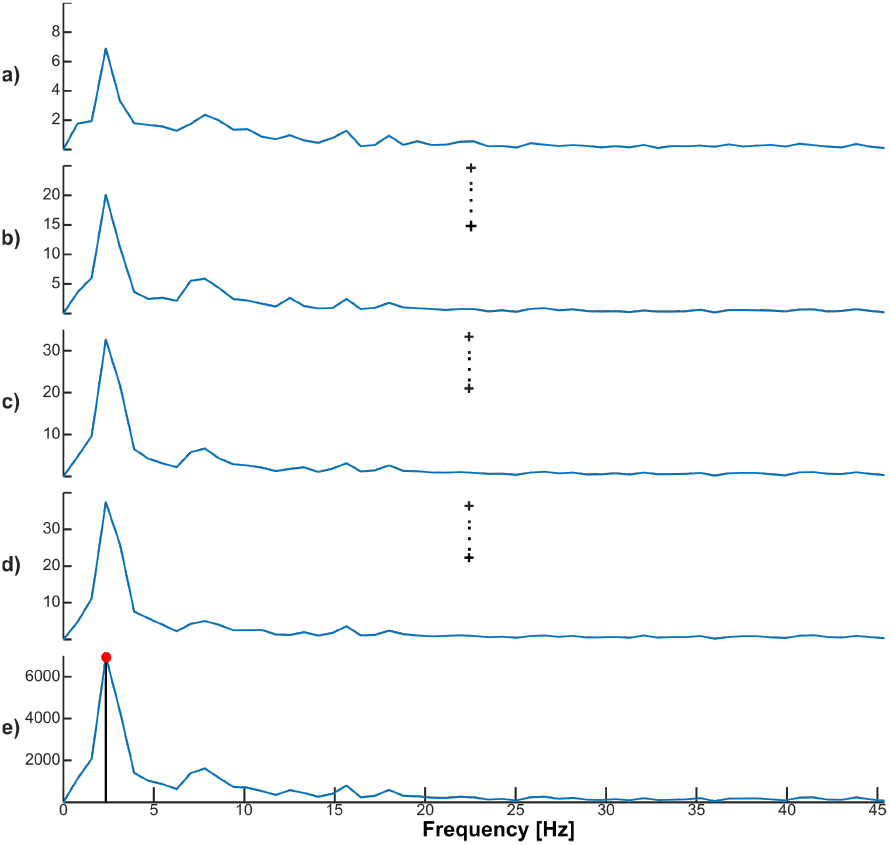
Graphic representation of the process for obtaining the beat frequency of a spermatozoon. a, b, c, d, and e) show the sum of the amplitude spectra for points 140, 240, 340 and 390, respectively (as a visual reference, these points are the same as those shown in Figure 5), for the two components (*Y, Z*) in the frequency domain single-sided plot. Plot e) displays the sum of the amplitude spectrum of all points, the red dot on the plot represents the maximum value of the spectrum to which the sperm beat frequency is associated.

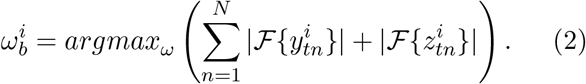

Finally, the asymmetry is taken as the mean of the eccentricity of the ellipses fitted to each spermatozoon in the head-fixed frame of reference. We performed the Wilcoxon Rank Sum Test to determine if there were differences between the two clusters for each measure described above (Figure 7), this test is used when the sizes of the samples are relatively small and they are not normally distributed, showing a significant difference.

**Figure 7:**
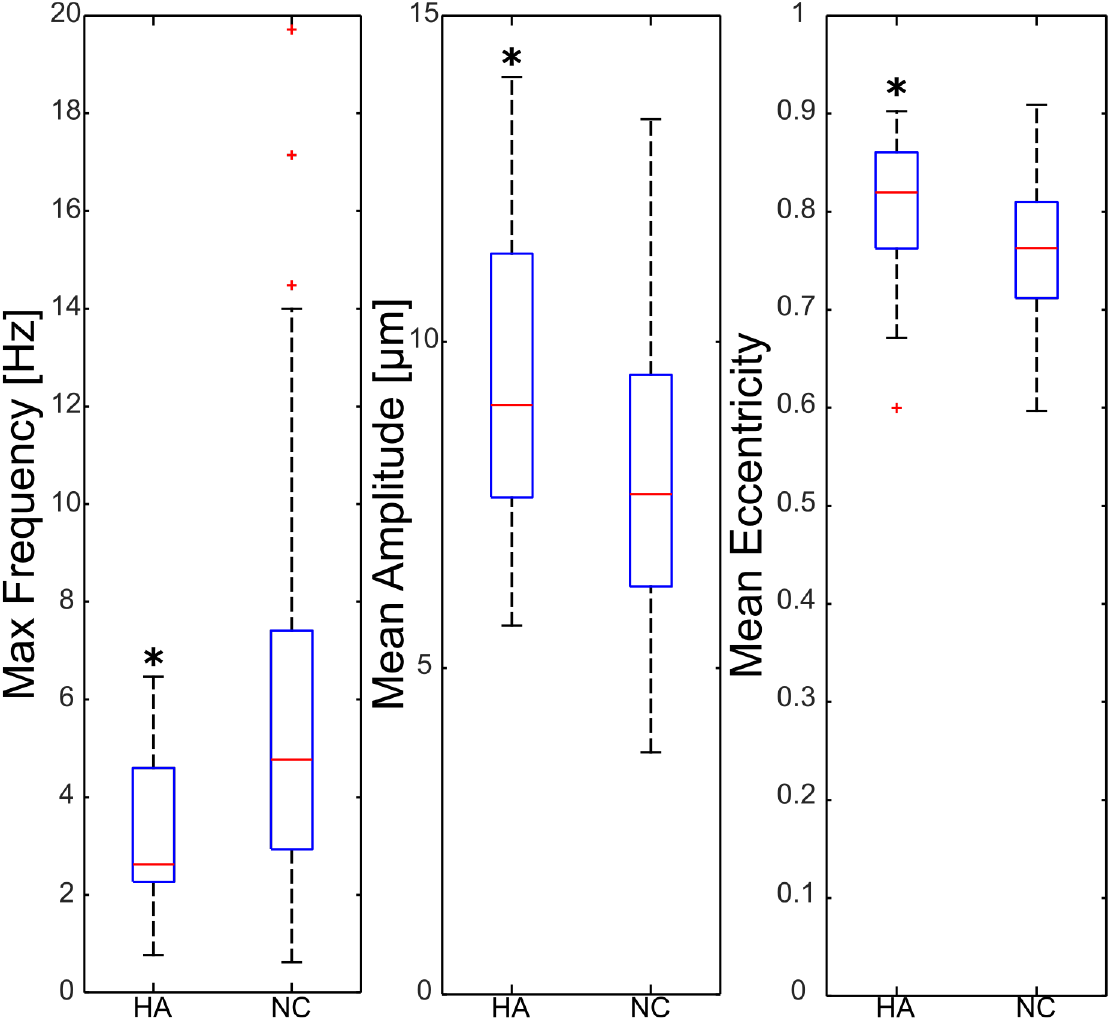
Boxplots of the frequency, amplitude, and asymmetry values for each motility group identified (hyperactivated and non-capacitated). The *p*−value for the Wilcoxon test for each feature is: 0.009, 0.01 and 0.0491 respectively ^∗^*P <* 0.05.

## 4 Conclusions and Discussion

The unsupervised classification of the flagellar beating of human spermatozoa into non-capacitated sperm, from those truly capacitated (hyperactivated) classes, was done using the proposed feature-based vector, which describes the average dynamics of the flagellar beat in 3D+t. It has been previously noted, for the 2D case, that hyperactivated spermatozoa have an asymmetrical beating pattern compared to non-capacitated ones [8]. Our proposed envelope of ellipses allows to generalize this description to the 3D case; since the ellipses with eccentricities close to one imply that elongated ellipses correspond to an asymmetric pattern while ellipses with eccentricities close to 0 are more symmetrical. In this sense, the 3D equivalent to the amplitude of the beat also increases in hyperactivated motility, this characteristic is obtained through the descriptor of the mean of the focus average of the ellipses. For the non-capacitated class, the mean of this feature is smaller than for the hyperactivated class. As expected, ≈ 17% of the induced to capacitate spermatozoa belong to the class truly capacitated, as mentioned in [9]. We emphasize that the entire set of non-capacitated sperm together with the failedto-capacitate from the induced to capacitate group were accurately classified in the green cluster. Since there is no ground-truth for 3D classification, we first attempted to compare the unsupervised classification results using the most common 2D characteristics of hyperactivation motility: frequency, amplitude, and asymmetry measured in 3D. This comparison shows that the frequency in the hyperactivated group is lower, while the mean amplitude is higher, and the asymmetry is greater than in non-capacitated group. The Wilcoxon test indicates that the two groups are statistically different in terms of each attribute (frequency, amplitude and asymmetry). Most results presented previously [25], coincide with those presented in this work, from what we can conclude that these descriptors reliably characterize the motility patterns. However, to provide a more accurate validation of the proposed feature-based descriptors for motility patterns, we are planning future work using fluorescent markers of hyperactivation. Finally, since the flagellar beating patterns have never been reported in 3D, our findings are encouraging. This approach could help to establish a criteria for classifying hyperactivated motility patterns in 3D+t by generalizing 2D descriptors, or by identifying unique characteristics that 3D+t data can provide.

## Acknowledgments

We thank Hermes Bloomfield-Gadêlha and Xiaomeng Ren members of the Polymaths Laboratory, Department of Engineering Mathematics, University of Bristol for their technical support.

## References

[1] S. T. Mortimer, G. Van der Horst, and D. Mortimer, “The future of computer-aided sperm analysis,” Asian journal of andrology, vol. 17, no. 4, p. 545, 2015.

[2] D. Bompart, A. García-Molina, A. Valverde, C. Caldeira, J. Yániz, M. N. de Murga, and C. Soler, “Casa-mot technology: how results are affected by the frame rate and counting chamber,” Reproduction, Fertility and Development, vol. 30, no. 6, pp. 810–819, 2018.

[3] C. Alquézar-Baeta, S. Gimeno-Martos, S. Miguel-Jiménez, P. Santolaria, J. Yániz, I. Palacín, A. Casao, J. Á. Cebrián-Pérez, T. Muiño-Blanco, and R. Pérez-Pé, “Opencasa: A new open-source and scalable tool for sperm quality analysis,” PLoS computational biology, vol. 15, no. 1, p. e1006691, 2019.

[4] R. Dcunha, R. S. Hussein, H. Ananda, S. Kumari, S. K. Adiga, N. Kannan, Y. Zhao, and G. Kalthur, “Current insights and latest updates in sperm motility and associated applications in assisted reproduction,” Reproductive Sciences, pp. 1–19, 2020.

[5] J.-w. Choi, L. Alkhoury, L. F. Urbano, P. Masson, M. VerMilyea, and M. Kam, “An assessment tool for computer-assisted semen analysis (casa) algrithms,” Scientific reports, vol. 12, no. 1, p. 16830, 2022.

[6] S. G. Goodson, S. White, A. M. Stevans, S. Bhat, C.-Y. Kao, S. Jaworski, T. R. Marlowe, M. Kohlmeier, L. McMillan, S. H. Zeisel, et al., “Casanova: a multiclass support vector machine model for the classification of human sperm motility patterns,” Biology of reproduction, vol. 97, no. 5, pp. 698–708, 2017.

[7] N. Sukcharoen, J. Keith, D. S. Irvine, and R. J. Aitken, “Definition of the optimal criteria for identifying hyperactivated human spermatozoa at 25 hz using in-vitro fertilization as a functional end-point,” Human Reproduction, vol. 10, no. 11, pp. 2928–2937, 1995.

[8] D. Waberski, S. S. Suarez, and H. Henning, “Assessment of sperm motility in livestock: Perspectives based on sperm swimming conditions in vivo,” Animal reproduction science, vol. 246, p. 106849, 2022.

[9] S. S. Suarez, “Control of hyperactivation in sperm,” Human reproduction update, vol. 14, no. 6, pp. 647–657, 2008.

[10] G. M. Centola, “Semen assessment,” Urologic Clinics, vol. 41, no. 1, pp. 163–167, 2014.

[11] S. Javadi and S. A. Mirroshandel, “A novel deep learning method for automatic assessment of human sperm images,” Computers in biology and medicine, vol. 109, pp. 182–194, 2019.

[12] P. Â. P. d. Silva, Supervised and unsupervised spermatozoa detection, classification and tracking in imaging data. PhD thesis, 2011.

[13] J. Riordon, C. McCallum, and D. Sinton, “Deep learning for the classification of human sperm,” Computers in biology and medicine, vol. 111, p. 103342, 2019.

[14] J. B. You, C. McCallum, Y. Wang, J. Riordon, R. Nosrati, and D. Sinton, “Machine learning for sperm selection,” Nature Reviews Urology, vol. 18, no. 7, pp. 387–403, 2021.

[15] B. J. Walker, S. Phuyal, K. Ishimoto, C.-K. Tung, and E. A. Gaffney, “Computer-assisted beat-pattern analysis and the flagellar waveforms of bovine spermatozoa,” Royal Society open science, vol. 7, no. 6, p. 200769, 2020.

[16] G. Corkidi, B. Taboada, C. Wood, A. Guerrero, and A. Darszon, “Tracking sperm in three-dimensions,” Biochemical and biophysical research communications, vol. 373, no. 1, pp. 125–129, 2008.

[17] G. Corkidi, F. Montoya, P. Hernández-Herrera, W. Ríos-Herrera, M. Müller, C. Treviño, and A. Darszon, “Are there intracellular ca2+ oscillations correlated with flagellar beating in human sperm? a three vs. two-dimensional analysis,” MHR: Basic science of reproductive medicine, vol. 23, no. 9, pp. 583–593, 2017.

[18] P. Hernandez-Herrera, F. Montoya, J. M. RendónMancha, A. Darszon, and G. Corkidi, “3-d +{t} human sperm flagellum tracing in low snr fluorescence images,” IEEE transactions on medical imaging, vol. 37, no. 10, pp. 2236–2247, 2018.

[19] H. Gadêlha, P. Hernández-Herrera, F. Montoya, A. Darszon, and G. Corkidi, “Human sperm uses asymmetric and anisotropic flagellar controls to regulate swimming symmetry and cell steering,” Science advances, vol. 6, no. 31, p. eaba5168, 2020.

[20] A. Gong, S. Rode, G. Gompper, U. B. Kaupp, J. Elgeti, B. Friedrich, and L. Alvarez, “Reconstruction of the three-dimensional beat pattern underlying swimming behaviors of sperm,” The European Physical Journal E, vol. 44, no. 7, p. 87, 2021.

[21] A. Danelakis, T. Theoharis, I. Pratikakis, and P. Perakis, “An effective methodology for dynamic 3d facial expression retrieval,” Pattern Recognition, vol. 52, pp. 174–185, 2016.

[22] M. Körtgen, G.-J. Park, M. Novotni, and R. Klein, “3d shape matching with 3d shape contexts,” in The 7th central European seminar on computer graphics, vol. 3, pp. 5–17, Budmerice Slovakia, 2003.

[23] P. Huang, J. Starck, and A. Hilton, “Temporal 3d shape matching,” in 4th European Conference on Visual Media Production, pp. 1–10, IET, 2007.

[24] A. K. Jain and R. C. Dubes, Algorithms for clustering data. Prentice-Hall, Inc., 1988.

[25] H. O. Hernández, P. Hernández-Herrera,F. Montoya, J. Olveres, H. Bloomfield-Gadêlha, A. Darszon, B. Escalante-Ramírez, and G. Corkidi, “3d+ t feature-based descriptor for unsupervised flagellar human sperm beat classification,” in 2022 44th Annual International Conference of the IEEE Engineering in Medicine & Biology Society (EMBC), pp. 488–492, IEEE, 2022.

[26] W. H. Organization et al., WHO laboratory manual for the examination and processing of human semen. World Health Organization, 2021.

[27] E. S. Maini, “Enhanced direct least square fitting of ellipses,” International Journal of Pattern Recognition and Artificial Intelligence, vol. 20, no. 06, pp. 939–953, 2006.

[28] E. De Lamirande, P. Leclerc, and C. Gagnon, “Capacitation as a regulatory event that primes spermatozoa for the acrosome reaction and fertilization.,” Molecular human reproduction, vol. 3, no. 3,pp. 175–194, 1997.

[29] V. Kay and L. Robertson, “Hyperactivated motility of human spermatozoa: a review of physiological function and application in assisted reproduction,” Human reproduction update, vol. 4, no. 6, pp. 776– 786, 1998.

[30] M. Zaferani, S. S. Suarez, and A. Abbaspourrad, “Mammalian sperm hyperactivation regulates navigation via physical boundaries and promotes pseudo-chemotaxis,” Proceedings of the National Academy of Sciences, vol. 118, no. 44, p. e2107500118, 2021.

[31] A. Agarwal, G. Virk, C. Ong, and S. S. Du Plessis, “Effect of oxidative stress on male reproduction,” The world journal of men’s health, vol. 32, no. 1, pp. 1–17, 2014.

[32] S. S. Suarez and A. Pacey, “Sperm transport in the female reproductive tract,” Human reproduction update, vol. 12, no. 1, pp. 23–37, 2006.

